# One drug multiple targets: An approach to predict drug efficacies on bacterial strains differing in membrane composition

**DOI:** 10.1101/423319

**Authors:** Ayan Majumder, Malay Ranjan Biswal, Meher K. Prakash

**Author notes:** Theoretical Science Unit, Jawaharlal Nehru Centre for Advanced Scientific Research, Jakkur, Bengaluru, India. Contributed equally to this work (Ayan Majumder, Malay Ranjan Biswal).

## Abstract

Rational design methodologies such as quantitative structure activity relationships (QSAR) have conventionally focused on screening through several drugs for their activity against a single target, either a bacterial protein or membrane. Recent concerns in drug design such as the development of drug resistance by membrane adaptation, or the undesirable damage to gut microbiota require a paradigm shift in activity prediction. A complementary approach capable of predicting the activity of a single drug against diverse targets, the diversity arising from bacterial adaptation or a heterogeneous composition with other helpful or harmful bacteria, is needed. As a first predictive step towards this goal, we develop a quantitative model for the activity of daptomycin on *Streptococcus aureus* strains with different membrane compositions, mainly varying in lysylation. The results of the predictions are good, and within the limits of the scarcely available data, hint at an interaction of daptomycin with the inner membrane. The complementary approach may in principle be extended to estimate the activity against gut bacterial membranes, when systematic data can be curated for training the model.

## Introduction

Antibiotic resistance is one of the major health threats in the decades to come. Rational antibiotic design methodologies use physico-chemical descriptors of drug candidates to predict their activity against well-defined targets, bacterial proteins or membranes. Many drug candidates are rejected for their toxicity, and others which translate as drugs become redundant with bacterial adaptation. In the last few decades, several pathogenic bacteria developed resistance to new antibiotics within a few years of their introduction. ^1^ Gut bacteria has been implicated in several important roles in human health and development^2^ and their disturbed balance can lead to seemingly unrelated disorders such as Parkinsons’.^3^ As such, the effects of commonly used drugs on gut bacteria^4^ are being evaluated. Only in the recent times physico-chemical rational intuitions are being used not just for efficacy prediction, but also to understand why most of the drug candidates do not reach advanced stages of clinical trials.^5^ The new generation of antibiotic design thus requires one to go beyond the traditional design paradigm and address a more comprehensive set of challenges rather than being an efficient drug against a well-defined target.

Computational methodologies such as quantitative structure activity relationships (QSAR) also were similarly motivated around designing drugs for well defined targets, by binding to the active site,^6,7^ or trapping reaction intermediates ^8^ or acting allosterically.^9^ When it was realized that bacteria develop resistance to these enzyme targeting drugs ^10,11^ easily, the focus of QSAR shifted towards evaluating the activity of cationic antimicrobial peptides (AMP) which act on bacterial membranes. The charge^12^ and amphipathic^13,14^ character of AMPs helps them disrupt the membrane, and a common bacterial adaptation is by lipid lysylation.^15^ Antimicrobial peptides which are part of the innate defense mechanisms^13,16–19^ continue to be the hope for treating several resistant strains of bacteria. But the focus ^20–22^ nevertheless remained on screening hundreds of drug candidates against a single target, which now needs to be complemented with the screening of drugs against multiple targets.

Pathogenic strains of *Streptococcus aureus* (*S. aureus*) which can cause severe skin and respiratory infections, develop resistance very fast to new antibiotics, and methicillin resistant *S. aureus* (MRSA) infections are especially problematic due to the lack of a suitable vaccine or antibiotic.^23,24^ Daptomycin is a cyclic lipopeptide antibiotic,^25–27^ it binds with the bacterial membrane in a Ca^2+^ dependent manner^28–32^ and the lipophilic acyl tail of daptomycin interacts with the membrane which then leads to K^+^ leakage and inhibition of protein, DNA, RNA synthesis.^33–35^ Daptomycin shows a significant activity against MRSA^36,37^ and vancomycin intermediate *S. aureus*.^28,38,39^ In this work, with the goal of broadening the scope beyond a conventional QSAR, we study the activity of daptomycin against strains of *S. aureus* characterized by different membrane compositions and adaptation.

## Methods

The experimental data on the membrane phospholipid composition of the different *S. aureus* strains, including methicillin resistant strain and their corresponding minimum inhibitory concentration (MIC) of daptomycin was curated from several published works.^40–51^ These works reported the total phospholipid composition - phosphatidylglycerol (PG), lysyl-PG (LPG) and cardiolipin (CL) - in the membrane bilayer (**Supplementary Table 1**). In addition, some works^44–51^ also reported the composition of the inner and outer leaflet LPG (referred to as iLPG and oLPG respectively) in the overall composition. The data was then used as follows to obtain the fractions of the phospholipids in the individual leaflets.

Assuming 2*N* lipid molecules in the bilayer, the composition was used to calculate the number of molecules (PG, LPG and CL) in the inner and outer leaflet. CL is assumed to be equally divided (%CL *× N*) between the two layers. (%iLPG*×*2*N*), (%PG*×* 2*N* +%oLPG*×*2*N* -%iLPG*×*2*N*)/4*N*, (%CL*× N*) are the number of LPG, PG and CL in the inner leaflet and (%oLPG*×*2*N*), (%PG*×* 2*N* + %iLPG *×* 2*N* - %oLPG*×*2*N*)/4*N*, (%CL *×N*) are the number of LPG, PG and CL in the outer leaflet. The percentage within each leaflet was then calculated based on these lipid molecule numbers. The phospholipid compositions thus derived for the inner and outer leaflets of the membrane, and the corresponding with daptomycin MIC values are given in **Supplementary Table 2**.

We have performed three separate artificial neural network calculations, using the 72 data points corresponding to total membrane composition, and 44 data points corresponding to the inner and outer leaflet compositions. In each of these cases, the data was randomly split into training, validation and test sets. Out of 72 data points with the total membrane composition data, 57 were chosen for training, 7 were chosen for validation and remaining 8 were chosen for testing. In the other two calculations using individual leaflet compositions from 44 membranes, we chose 35 data points for training, 4 for validation and 5 for testing.

Artificial neural network (ANN) model was used to obtain a relation between membrane composition and activity. The ANN model was based on scikit-learn,^52^ an open module for machine learning in Python. Logistic function was used as the activation function and low memory BFGS optimization algorithm was used as solver for this neural network.

With the total membrane composition data, 2500 trial runs were made, using 50 different randomized choices for the input biases in the neural network and 50 different randomized choices for the training and validation set. We used neural network models with an input, output and a hidden layer with 6, 8 or 10 neurons as an additional parameter. From all these trials, the best ANN models were selected by screening for two quality criteria, 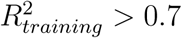 and 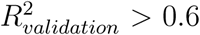. The same procedure was used when performing the calculations with the total membrane composition or with just the inner or outer leaflet compositions. One hidden layer of 8 neurons with an input and output layer gave the best result in our calculation for data corresponding total membrane composition and outer leaflet composition. On the other hand, for the inner leaflet, a hidden layer with 10 neurons gave the best results.

## Results

### Membrane descriptors

Driven by charges, daptomycin selectively attacks anionic the bacterial membranes, which are usually composed of phosphatidylglycerol (PG), cardiolipin (CL), lysyl-PG (LPG) and zwitterionic phosphatidylethanolamine (PE). The different strains of *S. aureus* which showed varying levels of drug resistance (MIC) were characterized by the compositions differing in PG, LPG and CL. We curated this data from 12 different studies^40–51^ (**Supplementary Table 1**). The curated data on average reflects that (**Supplementary Figure 1**) daptomycin binds and oligomerizes in the PG enriched region,^53^ and supports the intuition that the reduced toxicity to mammalian cells is due to their low PG content. CL^54^ and LPG^55^ on the other hand are negative factors decreasing the activity of daptomycin. However the exact relation is non-trivial (**Supplementary Figure 1**), since it depends on PG, LPG and CL varying simultaneously, and our goal in this work is to develop such relation.

### Membrane bilayer model

ANN model (**Methods** section) was used for obtaining a relation with MIC summarizing the activity of daptomycin on the strains characterized by different membrane compositions (**Methods**). The experimental MIC versus the MIC calculated using the total membrane composition of *S. aureus* is given in **Figure 1**. We obtained good results for the training 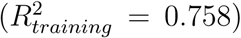, validation 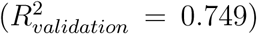 as well as for the test set predictions 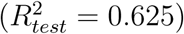.

**Figure 1:**
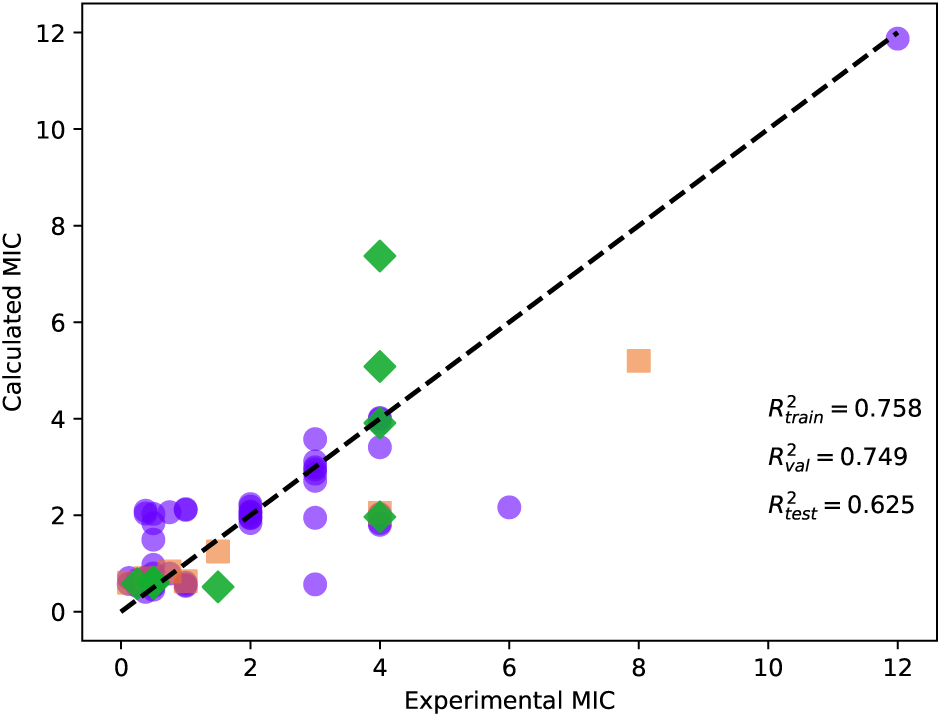
Comparison of the experimental MIC (*μ*g/mL) and MIC (*μ*g/mL) of daptomycin calculated using 8 neurons in the hidden layer on different overall membrane compositions. Training (purple circles), validation (orange squares) and test (green diamonds) sets are shown. The data used in the analysis is shown in **Supplementary Table 1**.

### Inner and outer leaflet models

In the curated data, the distribution of LPG between the inner and outer leaflets was available for 44 *S. aureus* strains. From this data, we derived the individual compositions for the two leaflets (**Methods section, Supplementary Table 2**). Using this derived information, we also performed two independent activity predictions considering only the inner and outer leaflet membrane compositions. The results are shown in **Figure 2**. The data analysis shows that the test predictions using only the inner leaflet 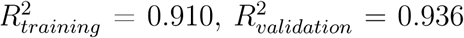 were better than those with the outer leaflet compositions 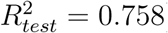.

**Figure 2:**
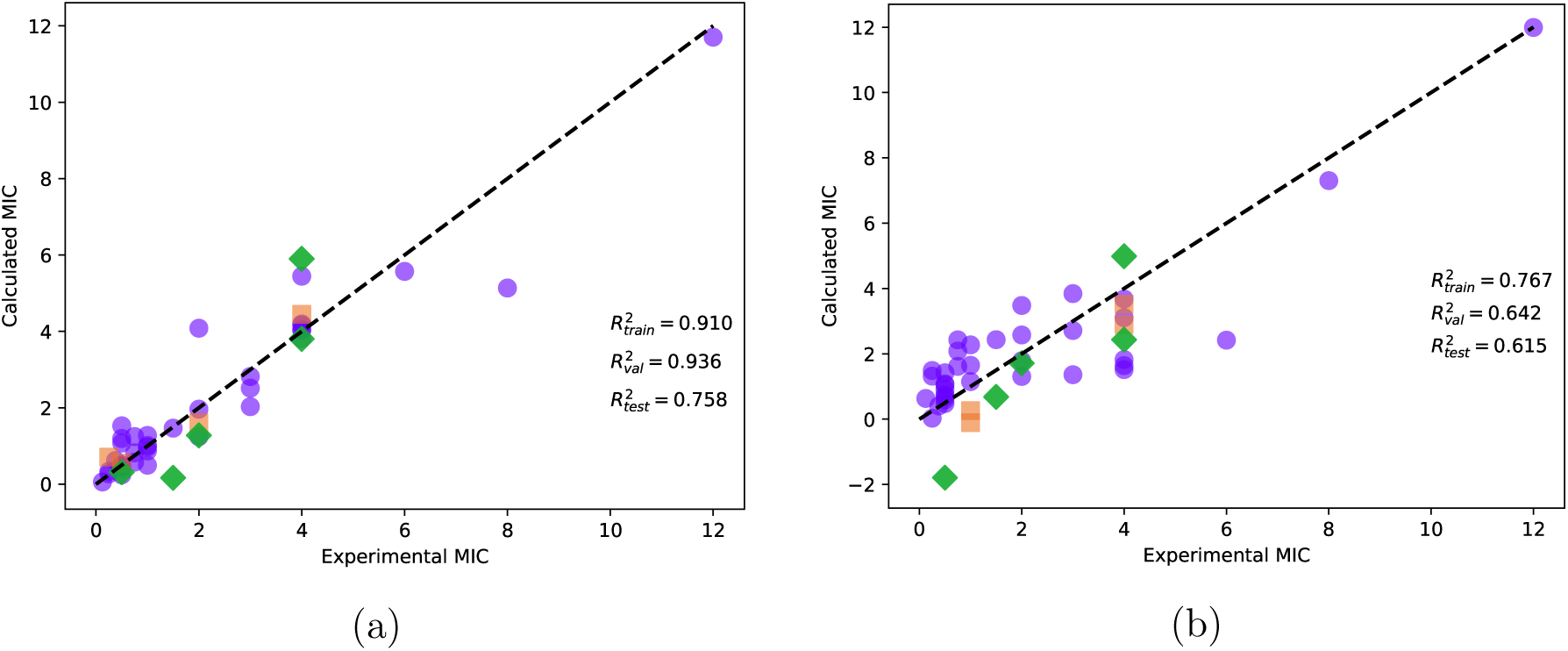
Comparison of the experimental (*μ*g/mL) and MIC (*μ*g/ml) of daptomycin calculated on different membrane compositions, using the data only from the a) inner leaflet, with 10 neurons in the hidden layer or b) the outer leaflet, with 10 neurons in the hidden layer. Training (purple circles), validation (orange squares) and test (green diamonds) sets are shown. The data used in the analysis is shown in **Supplementary Table 2.**

## Discussion

### One drug multiple membrane targets

In designing drugs to address antibiotic resistance, newer drugs such as antimicrobial peptide^56^ or their mimics^57^ which are cationic and effective against anionic bacterial membranes are being developed. As may be expected, bacteria adapt to such drugs, with a surface charge reduction by lysyl modification of the lipids. ^15^ However, rational design strategies, computational or experimental have focused mainly on designing the activity against a specific target, and it is not immediately apparent how effective the same drug remains when the bacteria adapts by lysylating a fraction of its PG. Procedurally our approach is the same as a standard QSAR, with a simple shift of focus from considering multiple drugs to multiple targets. We however believe that this simple change begins a new paradigm about quantitative structure activity predictions for the activity of a drug towards a broader range of targets, membranes in this case. Systematic experimental studies with a single drug against multiple targets have also been limited. To the best of our knowledge only on *S. aureus*, such data was available and was spread across several pieces of work. We curated the available data on daptomycin activity on *S. aureus* strains to develop a quantitative model. The model could be relevant for understanding how effective the drug remains as the composition of the membrane changes in serial passage experiments. ^44^ When systematically studied experimental data is available for training, similar approaches can be taken to study the effect of the same drug on membranes characterized by differences in lengths, types and saturation levels of lipids, potentially extendable to quantifying the unwanted effects of drugs on gut bacteria.

### Predicting daptomycin activity

Using ANN model we developed, we calculated the MIC of daptomycin for a systematic variation in the composition of the inner leaflet are show in **Figure 3** (and from overall and outer leaflet in **Supplementary Figure 2**). When the PG content is very high, MIC value is very low, which is in a good agreement with the previous experimental results. But when the PG concentration is below 50%, for a given value of LPG, MIC value decreases with increase in PG percentage and decrease in CL percentage. Interestingly, these non-monotonous trends are seen both in the experimental data (**Supplementary Figure 1**) as well as prediction from inner leaflet composition (**Figure 3**) This observation suggests that there are several ‘local equilibria’ in the membrane compositions that an adapting bacterium may find for improving its resistance (or increasing MIC), depending on the initial conditions or other constraints.

**Figure 3:**
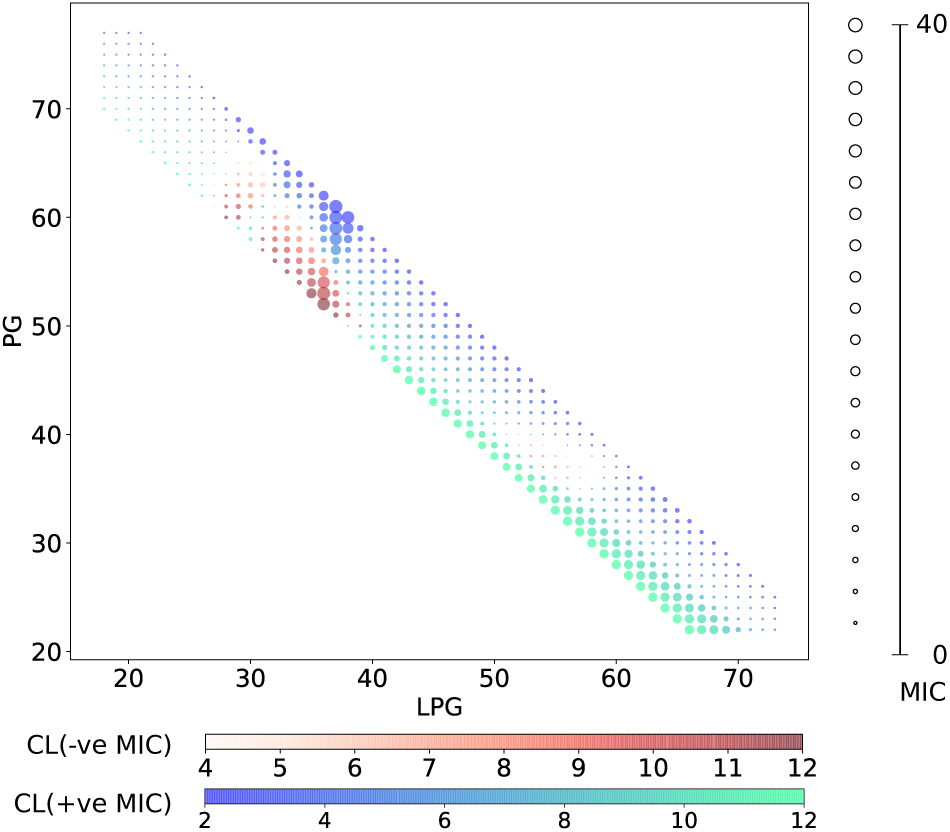
Change of daptomycin MIC (*μ*g/mL) values due to different PG and LPG concentrations in the inner leaflet of the membrane. The blue-green color represents the CL percentage for positive and the circle size represents MIC. Since we trained our model on the non-monotonous trends (**Supplementary Figure 1**), the model also resulted in a few negative MIC values (*μ*g/mL), represented in red color, over a small parametric region.

### Possible mechanism of action

Our calculations suggest that the composition of the inner leaflet better predicts the activity of daptomycin on *S. aureus*. Earlier experiments have suggested a correlation of the MIC with the LPG in the outer leaflet.^44^ However this correlation was based on few data points they obtained in that specific work, and the conclusions did not hold when the data from multiple studies were curated. As with the rest of the analysis in this article, the approach is conceptually new and the data available, especially for the individual leaflets, is limited. The conclusions need to be re-examined and adapted when more data becomes available.

## Conclusion

To the best of our knowledge a quantitative model that considers the effect of the same drug candidate on multiple membrane compositions has not been explored and it was developed in the present work to characterize the activity of daptomycin on different *S. aureus* strains. With the limited systematic data available, we could build a neural network based model which predicted the activity and suggested that the composition in the inner membrane is more critical than the overall or in the outer leaflets. The concepts can be strengthened and extended by modelling for *S. aureus* or for a heterogenous population when more systematic data becomes available for training and validation. While practically this one-drug-multitarget is a trivial extension of the concept of QSAR, conceptually it addresses an entirely new class of problems where the membrane adapts or the drug inadvertently acts on a heterogeneous population of bacteria including beneficial gut bacteria.

## Acknowledgment

We thank C. K. Sruthi for help with preparing the figures.

---

**Supplementary Table 1:** Total membrane phospholipid composition of the different *S. aureus* strain is given in terms of phosphatidylglycerol (PG), lysyl-PG (LPG), cardiolipin (CL), inner leaflet LPG (iLPG) and outer leaflet LPG (oLPG) and the daptomycin MIC values are given in *μ*g/mL. All the obtained experimental data are plotted in **Supplementary Figure 1**. Bacterial membrane phospholipid is mainly composed of anionic phosphatidylglycerol (PG), cardiolipin (CL) and cationic lysyl-phosphatidylglycerol (LPG).

| Strain | MIC | %LPG | %PG | %CL | %iLPG | %oLPG | References |
| --- | --- | --- | --- | --- | --- | --- | --- |
| C3 | 0.5 | 23.5 | 68.5 | 8 | - | - | 1 |
| C4 | 4 | 25.6 | 65.7 | 8.7 | - | - | 1 |
| C5 | 0.25 | 14.2 | 81.8 | 4 | - | - | 1 |
| C6 | 3 | 19.2 | 77.7 | 3 | - | - | 1 |
| C9 | 0.5 | 19.3 | 76 | 4.7 | - | - | 1 |
| C10 | 3 | 23.9 | 71.5 | 4.7 | - | - | 1 |
| C19 | 0.38 | 14.9 | 75.1 | 10 | - | - | 1 |
| C21 | 4 | 30.9 | 50.5 | 18.6 | - | - | 1 |
| C26 | 0.38 | 31.8 | 62.9 | 5.3 | - | - | 1 |
| C27 | 2 | 28.7 | 68.9 | 2.4 | - | - | 1 |
| C32 | 0.5 | 21.9 | 71.2 | 6.9 | - | - | 1 |
| C33 | 2 | 24.1 | 73.1 | 2.7 | - | - | 1 |
| C36 | 0.5 | 15.1 | 81.5 | 3.4 | - | - | 1 |
| C37 | 3 | 26.1 | 69 | 5 | - | - | 1 |
| C40 | 0.25 | 26 | 68.4 | 5.6 | - | - | 1 |
| C41 | 3 | 30 | 59.7 | 10.3 | - | - | 1 |
| CB1118<sup>1</sup> | 1 | 12.37 | 83.96 | 5.38 | 11.16 | 1.21 | 2 |
| CB2201 | 1.5 | 12.29 | 86.37 | 3.6 | 10.92 | 1.37 | 2 |
| CB2202 | 3 | 13.34 | 80.95 | 8.59 | 11.64 | 1.7 | 2 |
| CB2203 | 6 | 17.96 | 72.93 | 12.7 | 15.41 | 2.56 | 2 |
| CB2205 | 12 | 24.61 | 70.02 | 7.43 | 18.61 | 6 | 2 |

<sup>1</sup>CB1118 strain was used as a parental strain in two independent serial passage studies<sup>2</sup> and.<sup>3</sup> As noted in the latter study, the novel mutations evolved in the latter study and led to two notable changes - a decreased LPG after certain time and an increased production of carotenoid. Since the influence of carotenoids was an additional parameter for which no other study we considered provided the data, we excluded the latter study from our analysis.

**Supplementary Table 1:** Total membrane phospholipid composition of the different *S. aureus* strain is given in terms of phosphatidylglycerol (PG), lysyl-PG (LPG), cardiolipin (CL), inner leaflet LPG (iLPG) and outer leaflet LPG (oLPG) and the daptomycin MIC values are given in $\mu\text{g/mL}$ .
| Strain | MIC | %LPG | %PG | %CL | %iLPG | %oLPG | References |
| --- | --- | --- | --- | --- | --- | --- | --- |
| 1A | 0.25 | 13 | 81 | 6 | 11 | 2 | 4,5 |
| 1C | 4 | 26 | 68 | 6 | 22 | 5 | 4,5 |
| 2A | 0.5 | 11 | 84 | 5 | 9 | 2 | 4,5 |
| 2C | 2 | 24 | 67 | 9 | 19 | 4 | 4,5 |
| 3A | 0.5 | 22 | 69 | 9 | 19 | 3 | 4,5 |
| 3B | 4 | 16 | 80 | 4 | 13 | 3 | 4,5 |
| CB1483 | 0.25 | 15.3 | 77.21 | 7.49 | 13.39 | 1.91 | 6 |
| CB185 | 4 | 35.96 | 52.31 | 11.73 | 32.1 | 3.85 | 6 |
| CB5079 | 0.5 | 15.43 | 72.12 | 12.44 | 13.63 | 1.8 | 6 |
| CB5080 | 2 | 26.18 | 64.08 | 9.75 | 24.78 | 1.39 | 6 |
| CB5083 | 0.25 | 12.07 | 83.3 | 4.63 | 10.15 | 1.92 | 6 |
| CB5082 | 4 | 21.61 | 73.58 | 4.82 | 19.24 | 2.37 | 6 |
| CB5088 | 0.5 | 15.36 | 77.66 | 6.97 | 13.61 | 1.75 | 6 |
| CB5089 | 2-4 | 24.91 | 65.93 | 9.16 | 22.62 | 2.29 | 6 |
| CB1631 | 0.5 | 12.08 | 80.41 | 7.51 | 10.25 | 1.83 | 6 |
| CB1634 | 4 | 20.61 | 71.75 | 7.63 | 18.68 | 1.93 | 6 |
| CB1663 | 0.5 | 11.96 | 83.2 | 4.84 | 10.24 | 1.72 | 6 |
| CB1664 | 4 | 16.05 | 81.33 | 2.62 | 14.69 | 1.36 | 6 |
| CB5057 | 0.5 | 15.91 | 79.25 | 4.85 | 14.32 | 1.59 | 6 |
| CB5059 | 4 | 27.22 | 69.92 | 2.87 | 24.77 | 2.44 | 6 |
| CB5062 | 0.5 | 12.71 | 79.9 | 7.4 | 11.49 | 1.21 | 6 |
| CB5063 | 8 | 31.55 | 59.01 | 9.44 | 29.23 | 2.33 | 6 |
| CB5015 | 1 | 14.06 | 83.31 | 2.63 | 12.73 | 1.32 | 6 |
| CB5016 | 4 | 19.27 | 77.2 | 3.53 | 17.61 | 1.66 | 6 |
| C11 | 0.38 | 18 | 78 | 5 | - | - | 7 |
| C12 | 3 | 25 | 68 | 7 | - | - | 7 |
| C28 | 0.12 | 15 | 79 | 6 | - | - | 7 |
| C29 | 2 | 23 | 70 | 7 | - | - | 7 |
| C44 | 0.38 | 20 | 74 | 5 | - | - | 7 |
| C45 | 4 | 26 | 68 | 6 | - | - | 7 |
| Newman | 0.5 | 17 | 72 | 11 | - | - | 8 |
| Newman mprF | 0.125 | 0 | 86 | 14 | - | - | 8 |
| CS295L | 1 | 12 | 82 | 7 | - | - | 8 |
| CS295L+L826F | 0.38 | 0 | 91 | 9 | - | - | 8 |
| CT345A | 2 | 18 | 72 | 10 | - | - | 8 |
| CT345A+L826F | 0.38 | 0 | 91 | 9 | - | - | 8 |
| None | 0.5 | 13.2 | 81.5 | 5.3 | 11.2 | 2 | 9 |
| None | 1 | 15.9 | 75.5 | 8.7 | 13.6 | 2.3 | 9 |
| None | 2 | 24.5 | 68 | 7.5 | 22 | 2.5 | 9 |
| L271 | 0.125 | 14.35 | 83.05 | 2.59 | 13.28 | 1.07 | 10 |
| L8 | 2 | 31.75 | 66.78 | 1.47 | 30.86 | 0.89 | 10 |
| L16 | 0.75 | 31.01 | 66.12 | 2.87 | 29.55 | 1.46 | 10 |
| L56 | 2 | 39.15 | 58.6 | 2.26 | 36.27 | 2.88 | 10 |
| L76 | 0.38 | 16.81 | 72.27 | 11 | 14.76 | 2.05 | 10 |
| SA144 | 0.75 | 17.45 | 74.28 | 8.27 | 15.38 | 2.08 | 11 |
| SA145 | 0.75 | 17.86 | 69.92 | 12.22 | 15.24 | 2.63 | 11 |
| SA147 | 1.5 | 18.86 | 69.42 | 11.72 | 15.04 | 3.83 | 11 |
| MRSA 11/11 | 1 | 18.91 | 75.24 | 5.86 | 17.17 | 1.73 | 12,13 |
| MRSA 11/17 | 1 | 19.63 | 72.57 | 7.81 | 18.16 | 1.46 | 12,13 |
| MRSA 11/21 | 3 | 34.16 | 62.45 | 5.68 | 32.15 | 2.72 | 12,13 |
| REF2145 | 4 | 36.66 | 55.25 | 7.8 | 34.67 | 2.29 | 12,13 |

**Supplementary Figure 1:**
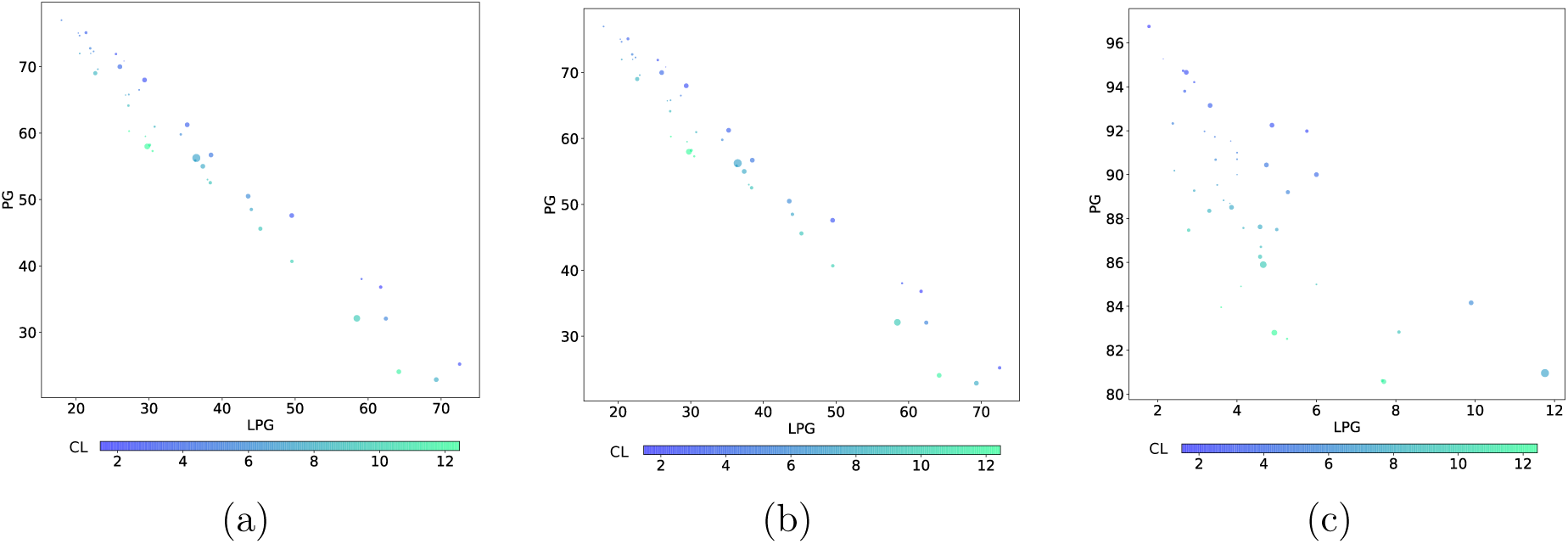
Change in daptomycin MIC values with different a) total membrane, b) inner leaflet and c) outer leaflet phospholipid compositions obtained from experimental data. The radius of the circle represents MIC (*μ*g/ml) and the change in color represents CL concentration.

**Supplementary Table 2:**
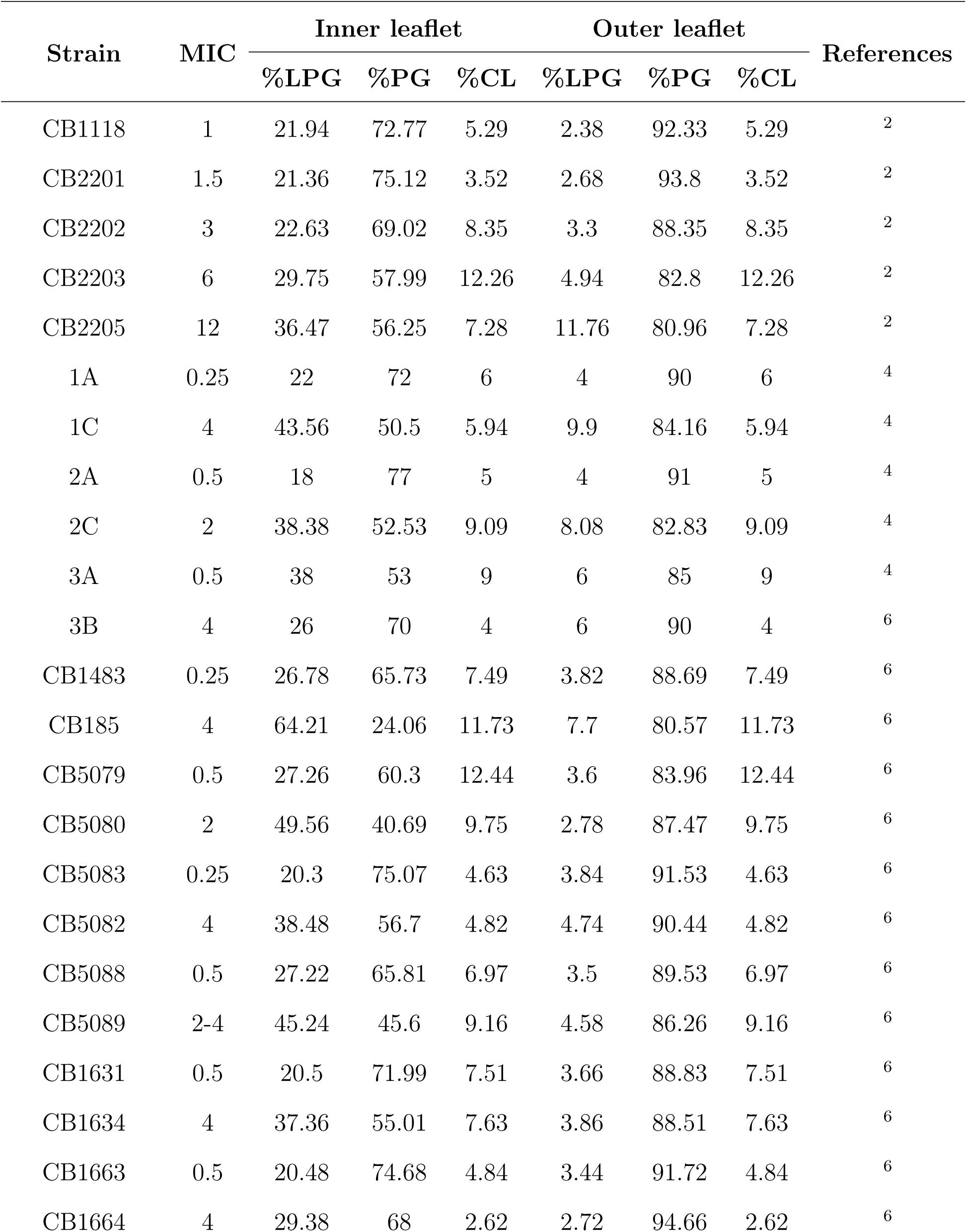
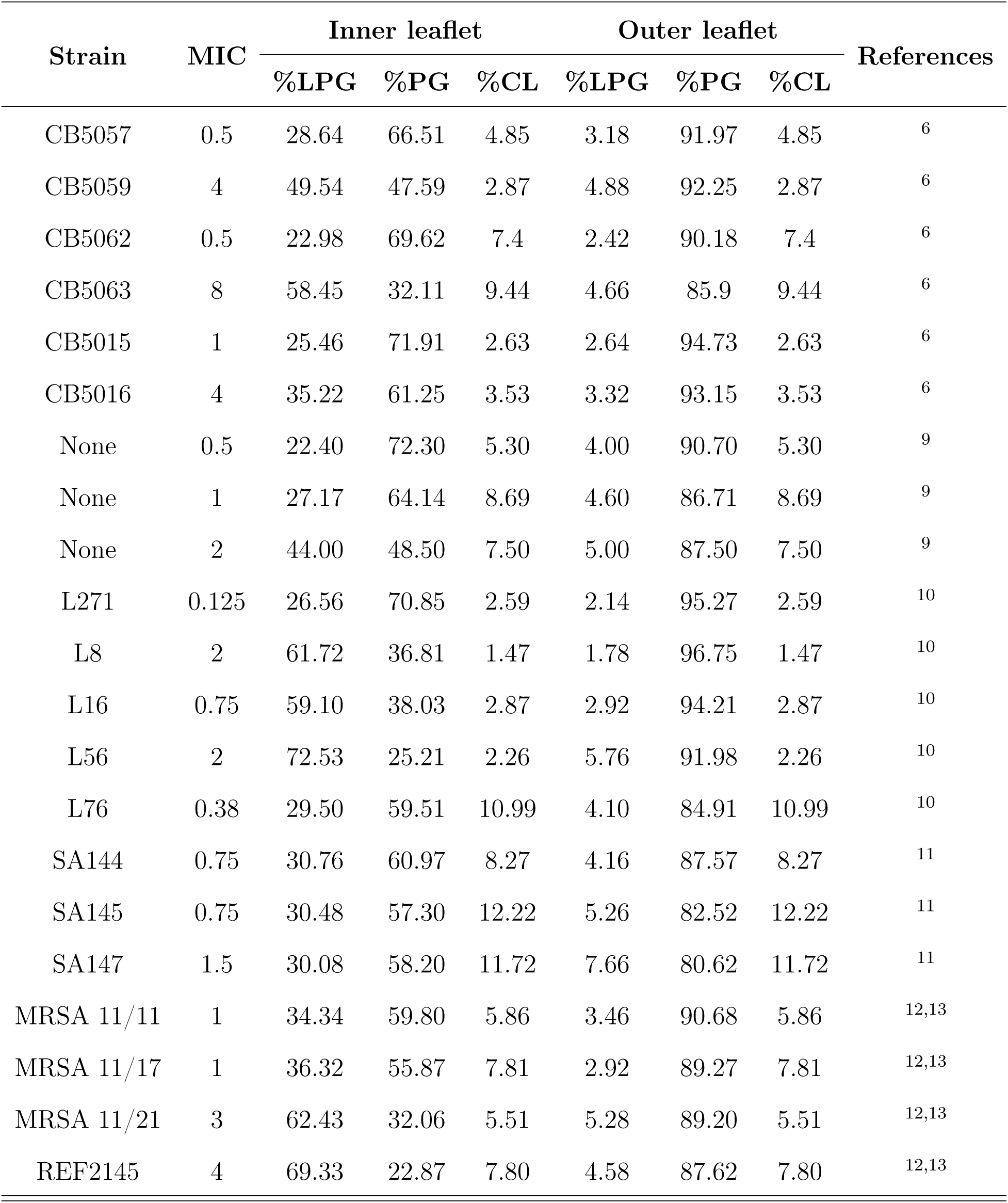
Inner and outer leaflet of membrane phospholipid composition of the different *S. aureus* strains in terms of lyslphosphatidylglycerol (lysl-PG), phosphatidylglycerol (PG), cardiolipin (CL) and the daptomycin MIC values in *μ*g/ml. Although the calculation shows that the daptomycin activity can be predicted well from the inner leaflet composition, but the total membrane and outer leaflet composition can also predict the daptomycin activity to some extent. The prediction for the possible membrane composition on the basis of total and outer leaflet compositions are shown in **Supplementary Figure 2**.

| Strain | MIC | Inner leaflet |  |  | Outer leaflet |  |  | References |
| --- | --- | --- | --- | --- | --- | --- | --- | --- |
|  |  | %LPG | %PG | %CL | %LPG | %PG | %CL |  |
| CB1118 | 1 | 21.94 | 72.77 | 5.29 | 2.38 | 92.33 | 5.29 | 2 |
| CB2201 | 1.5 | 21.36 | 75.12 | 3.52 | 2.68 | 93.8 | 3.52 | 2 |
| CB2202 | 3 | 22.63 | 69.02 | 8.35 | 3.3 | 88.35 | 8.35 | 2 |
| CB2203 | 6 | 29.75 | 57.99 | 12.26 | 4.94 | 82.8 | 12.26 | 2 |
| CB2205 | 12 | 36.47 | 56.25 | 7.28 | 11.76 | 80.96 | 7.28 | 2 |
| 1A | 0.25 | 22 | 72 | 6 | 4 | 90 | 6 | 4 |
| 1C | 4 | 43.56 | 50.5 | 5.94 | 9.9 | 84.16 | 5.94 | 4 |
| 2A | 0.5 | 18 | 77 | 5 | 4 | 91 | 5 | 4 |
| 2C | 2 | 38.38 | 52.53 | 9.09 | 8.08 | 82.83 | 9.09 | 4 |
| 3A | 0.5 | 38 | 53 | 9 | 6 | 85 | 9 | 4 |
| 3B | 4 | 26 | 70 | 4 | 6 | 90 | 4 | 6 |
| CB1483 | 0.25 | 26.78 | 65.73 | 7.49 | 3.82 | 88.69 | 7.49 | 6 |
| CB185 | 4 | 64.21 | 24.06 | 11.73 | 7.7 | 80.57 | 11.73 | 6 |
| CB5079 | 0.5 | 27.26 | 60.3 | 12.44 | 3.6 | 83.96 | 12.44 | 6 |
| CB5080 | 2 | 49.56 | 40.69 | 9.75 | 2.78 | 87.47 | 9.75 | 6 |
| CB5083 | 0.25 | 20.3 | 75.07 | 4.63 | 3.84 | 91.53 | 4.63 | 6 |
| CB5082 | 4 | 38.48 | 56.7 | 4.82 | 4.74 | 90.44 | 4.82 | 6 |
| CB5088 | 0.5 | 27.22 | 65.81 | 6.97 | 3.5 | 89.53 | 6.97 | 6 |
| CB5089 | 2-4 | 45.24 | 45.6 | 9.16 | 4.58 | 86.26 | 9.16 | 6 |
| CB1631 | 0.5 | 20.5 | 71.99 | 7.51 | 3.66 | 88.83 | 7.51 | 6 |
| CB1634 | 4 | 37.36 | 55.01 | 7.63 | 3.86 | 88.51 | 7.63 | 6 |
| CB1663 | 0.5 | 20.48 | 74.68 | 4.84 | 3.44 | 91.72 | 4.84 | 6 |
| CB1664 | 4 | 29.38 | 68 | 2.62 | 2.72 | 94.66 | 2.62 | 6 |
|  |  | %LPG | %PG | %CL | %LPG | %PG | %CL |  |
| CB5057 | 0.5 | 28.64 | 66.51 | 4.85 | 3.18 | 91.97 | 4.85 | 6 |
| CB5059 | 4 | 49.54 | 47.59 | 2.87 | 4.88 | 92.25 | 2.87 | 6 |
| CB5062 | 0.5 | 22.98 | 69.62 | 7.4 | 2.42 | 90.18 | 7.4 | 6 |
| CB5063 | 8 | 58.45 | 32.11 | 9.44 | 4.66 | 85.9 | 9.44 | 6 |
| CB5015 | 1 | 25.46 | 71.91 | 2.63 | 2.64 | 94.73 | 2.63 | 6 |
| CB5016 | 4 | 35.22 | 61.25 | 3.53 | 3.32 | 93.15 | 3.53 | 6 |
| None | 0.5 | 22.40 | 72.30 | 5.30 | 4.00 | 90.70 | 5.30 | 9 |
| None | 1 | 27.17 | 64.14 | 8.69 | 4.60 | 86.71 | 8.69 | 9 |
| None | 2 | 44.00 | 48.50 | 7.50 | 5.00 | 87.50 | 7.50 | 9 |
| L271 | 0.125 | 26.56 | 70.85 | 2.59 | 2.14 | 95.27 | 2.59 | 10 |
| L8 | 2 | 61.72 | 36.81 | 1.47 | 1.78 | 96.75 | 1.47 | 10 |
| L16 | 0.75 | 59.10 | 38.03 | 2.87 | 2.92 | 94.21 | 2.87 | 10 |
| L56 | 2 | 72.53 | 25.21 | 2.26 | 5.76 | 91.98 | 2.26 | 10 |
| L76 | 0.38 | 29.50 | 59.51 | 10.99 | 4.10 | 84.91 | 10.99 | 10 |
| SA144 | 0.75 | 30.76 | 60.97 | 8.27 | 4.16 | 87.57 | 8.27 | 11 |
| SA145 | 0.75 | 30.48 | 57.30 | 12.22 | 5.26 | 82.52 | 12.22 | 11 |
| SA147 | 1.5 | 30.08 | 58.20 | 11.72 | 7.66 | 80.62 | 11.72 | 11 |
| MRSA 11/11 | 1 | 34.34 | 59.80 | 5.86 | 3.46 | 90.68 | 5.86 | 12,13 |
| MRSA 11/17 | 1 | 36.32 | 55.87 | 7.81 | 2.92 | 89.27 | 7.81 | 12,13 |
| MRSA 11/21 | 3 | 62.43 | 32.06 | 5.51 | 5.28 | 89.20 | 5.51 | 12,13 |
| REF2145 | 4 | 69.33 | 22.87 | 7.80 | 4.58 | 87.62 | 7.80 | 12,13 |

**Supplementary Figure 2:**
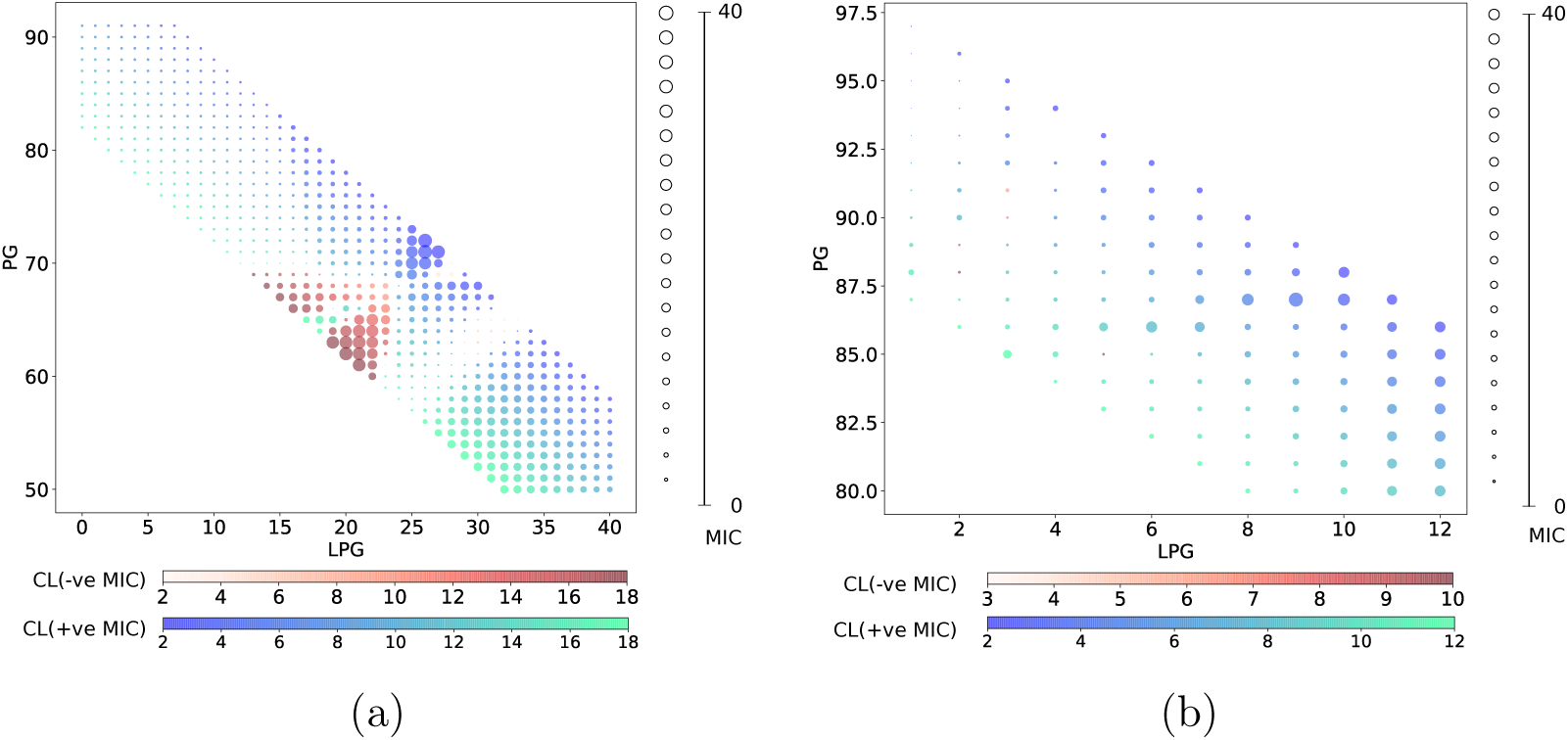
Change of daptomycin MIC (*μ*g/ml) values due to different PG and LPG concentration of the total membrane. The blue-green and red color represents the change in CL percentage for postive and negative MIC values respectively and radius of the circle increases with increase in absolute value of MIC.

